# VCF2Prot: An Efficient and Parallel Tool for Generating Personalized Proteomes from VCF Files

**DOI:** 10.1101/2022.01.21.477084

**Authors:** Hesham ElAbd, Frauke Degenhardt, Tobias L. Lenz, Andre Franke, Mareike Wendorff

**Affiliations:** Institute of Clinical Molecular Biology, Christian-Albrechts-University of Kiel, Kiel, Germany; Research Unit for Evolutionary Immunogenomics, Department of Biology, University of Hamburg, Hamburg, Germany

**Author notes:** joint coordination and supervision of project (Andre Franke, Mareike Wendorff). to whom correspondence shall be addressed /.

## Abstract

**Motivation:** The ability to generate sample-specific protein sequences is a crucial step in neo-antigen discovery, cancer vaccine development, and proteogenomics. The revolutionary increase in the throughput of sequencers has fueled large-scale genomic and transcriptomic studies, holding great promises for the emerging field of personalized medicine. However, most sequencing projects store their sequencing data in an abbreviated variant calling format (VCF) that is not immediately amenable to subsequent proteomic and peptidomic analyses. Furthermore, data processing of such increasingly massive genome-scale datasets calls for parallel and concurrent programming, and consequently refactoring of existing algorithms and/or the development of new parallel algorithms.

**Results:** Here, we introduce sequence intermediate representation (SIR), a novel and generic algorithm for generating personalized or sample-specific protein sequences from a consequence-called VCF file and the corresponding reference proteome. An implementation of SIR, named VCF2Prot, was developed to aid personalized medicine and proteogenomics by generating personalized proteomes in FASTA format from a collection of consequence-called genomic alterations stored in a VCF file. Benchmarking VCF2Prot against the recently published PrecisionProDB showed an ~1000-fold improvement in runtime (depending on the input size). Furthermore, in a scale-up study VCF2Prot processed a VCF file containing 99,254 variants observed across 8,192 patients in ~ 11 minutes, demonstrating the massive improvement in the execution speed and the utility of SIR and VCF2prot in bridging large-scale genomic and proteomic studies.

**Availability and Implementation:** *VCF2Prot* comes with a permissive MIT-license, enabling the commercial and non-commercial utilization of the tool. The source code along with precompiled versions for Linux/Mac OS are available at https://github.com/ikmb/vcf2prot. The modular units used for building *VCF2Prot* are available as a Rust crate at https://crates.io/crates/ppgg with documentations and examples at https://docs.rs/ppgg/0.1.4/ppgg/ under the same MIT-license.

## Introduction

Generating sample-specific proteomes is a fundamental step in proteogenomics and cancer neoantigen discovery. In the field of proteogenomics, next-generation sequencing is commonly used to generate a sample-specific protein sequence database for identifying the spectra obtained through mass spectrometers [1]. Similarly, in neoantigen discovery, generating sample-specific protein sequences is needed to identify mutated peptides that might generate an immune response towards cancer [2]. Furthermore, protein and peptide sequences are the main input for predicting peptide-HLA interactions [3, 4] and are a corner-stone for unraveling functional HLA-associations in diseases using peptidome-wide association studies [5].

Different tools have been developed to generate personalized proteomes from sequencing data, for examples, *PrecisionProDB* [6], *Pypgatk* [7], and *CustomProDB* [8]. These tools differ in their required input format, the execution logic, the programing language they are implemented in and the output format they generate. *CustomProDB* [8] is an R-package specialized in generating personalized protein sequences from RNA-Seq data, but not from genome sequence data. *Pypgatk* [7] is a python library for generating personalized protein databases from alterations in canonical and non-canonical genes, but it only considers single nucleotide polymorphisms and no structural variants. *PrecisionProDB* [6] is a recently published tool implemented in Python for generating sample-specific protein sequences from genomic alterations. However, it can only handle one sample at a time and hence it cannot be used for large scale applications where, thousands of samples need to be analyzed.

With the recent increase in sequencing throughput, decrease in cost, and improvement in mass spectrometry sensitivity, omics data production is continuously growing. Hence, tools and algorithms for handling large scale sequencing datasets are urgently needed. To address this problem, we first developed sequence **i**ntermediate **r**epresentation (SIR), a new inherently parallel algorithm for generating personalized protein sequences from genomic alterations. Secondly, we developed *VCF2Prot*, a freely available command line tool implemented in the Rust programming language that provides an optimized implementation of SIR.

## Material and Methods

### I Problem Formulation and Algorithm Description

We can define any personalized sequence, *S*_*new*_, as the concatenation of a collection of subsequences, or a collection of blocks, as described in Eq.1.

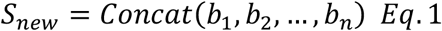

where *b*_1_, … … *b*_*n*_ represent a collection of subsequences. These blocks are obtained from two sources: the first is sequences copied from the reference proteome (for stretches of sequence without individual alterations) and the second is the alternative sequences due to sample-specific genomic alterations (**Table S1 & Fig. S1**). Thus, the first step in generating personalized sequences is to identify contingent blocks of subsequences from the reference and the altered stream. To do this, the position relative to the reference, and the type of alteration must be identified. By using sequencing-technologies, e.g. exome-sequencing, and variant calling algorithms, e.g. *BCFtools/csq* [9] (**Table S1**), the position and type of coding variants can be obtained proteome-wide. Next, the position and type of alteration are translated into a representation that describes the generation of sequence blocks, referred to hereafter as *Instruction. Instructions* act as intermediate, fixed-size, and uniform representation for a genomic alteration. It is composite of 5 main components, namely, code which is a one-character identity for the alteration type (**Table S2**), position in the reference, position in the altered sequence, pointer to a char array storing the altered sequence and a number storing the length of alterations.

Besides acting as a simplified internal representation, *Instructions* enable semantic equivalence where different genetic mutations can have an identical effect on the protein level, i.e. they result in the same protein end product. For example, a frameshift mutation that causes a premature termination of protein translation is synonymous with stop-gain mutation. Hence, different *Instructions* can be casted into different types, i.e. *Instructions* with different instruction-code and, thus, simplifying the problem for down-stream code. Once instructions have been created, they are validated and translated into an even more simplified representation referred to as *Tasks* and a corresponding string of resulting sequences. *Tasks* are the simplest representation used by the algorithm. Structurally, a *Task* is a tuple t composed of four elements, first, the stream code *c*, which describes the source of reading data, e.g., reference or alteration. Second, the position *p*, which describes the start of the block in the input stream. Third, the length *l*, which describes the length of the block and finally, the position in the new sequence, *r* (**Fig. 1**).

**Figure 1:**
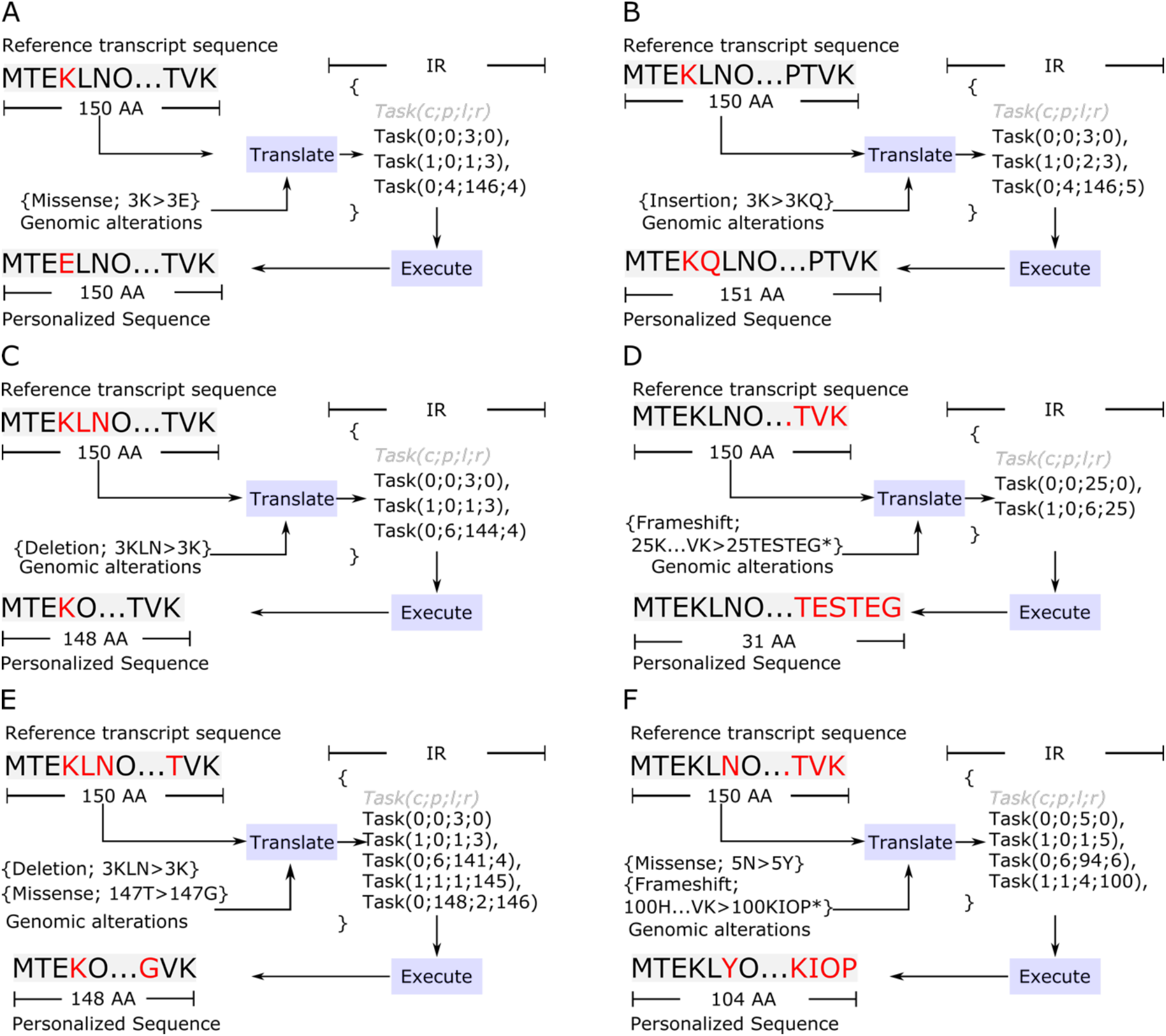
A schematic summary for the translation of different genomic alterations as given by *BCFTools/csq* [9] (changed to be zero-indexed) into the intermediate representation (IR). **A** shows the translation of a missense mutation where the amino acid K is replaced with E. The resulting IR consists therefore of 3 tasks: 1. Take the first three amino acids from the reference. 2. Take the alternate from the genomic alteration. 3. Take the reference from position 4 till the end. **B** the translation of an in-frame insertion with the amino acid Q after the 4^th^ amino acid K. **C** shows the translation of a deletion after the 4^th^ amino acid where the amino acids KLN are replaced by only the K. **D** shows the translation of a frameshift mutation where the subsequence starting from position 25 until the end the of the protein is replaced with the sequence TESTEG. **E** shows the translation of a more complex case where two alterations are observed on the protein, the first is a deletion at the 3^rd^ amino acid and the second is a missense mutation of the 148^th^ amino acid. **F** shows the translation of two alterations: a missense in the 5^th^ position and a frameshift at the 100^th^ amino acids. All indices in the figure are zero-based to reflect the same indexing system used by VCF2Prot’s Rust-based implementation.

A collection of *Tasks* describing the generation of a collection of sequence blocks is referred to hereafter as the intermediate representation (IR) of the sequence (**Fig. 1 A-B**). **Fig. S1** describes the translation of a reference sequence and an observed alteration to an IR. First, for a given collection of alterations, a collection of *Tasks* is created, through the generation of *Instructions*. Once the alterations in each transcript, i.e. genomic alteration in the protein sequence of a transcript, have been projected into the IR space, they can be combined, concatenated, and shifted, i.e. moved along the indexing axis. This can be exploited to concatenate the IR of each sequence to generate a personalized proteome-wide IR.

As shown in **Fig. S1**, to generate a proteome-wide IR, first, alterations are grouped by transcript, i.e. all alterations that occurred in the protein sequence of one transcript are grouped together. Second, each sequence is projected into the IR space. Third, the IR of each sequence is concatenated and reindexed to generate a proteome-wide representation. As seen in **Fig. S1**, each sample is processed independently from other samples, enabling different samples to be processed in parallel by different threads.

### II Inspection and Validation of the Translation into Instructions

As stated above, *Instructions* are the first step toward generating an IR and hence the validity of translations is of paramount importance to ensure the correct generation of IR. Hence, three tests were implemented to validate the correctness of input mutations and of translations, first, position uniqueness, second isolated boundaries, and third prohibited sequences. Position-uniqueness refers to the requirement that each instruction must have a unique starting position in the reference and the altered sequence and hence having two mutations at the same position is prohibited. “Isolated boundaries” refers to the requirement that mutations should not overlap. Finally, “prohibited sequences” refers to an ‘illegal’ sequence of mutations in a transcript, for example, after a frameshift, stop-gain, or a stop-loss no independent mutation is allowed. If any of the three tests fail, the input collection of mutation and the translation is considered invalid and hence, it is by default ignored, i.e., skipped, and an error message is printed to the user.

Many reasons can cause such translation failures, for example, software bugs in the encoding, parsing, and decoding of the bitmask, errors in the consequence calling algorithm due to the complex genetic architecture of the sample, or due to mismatched references where the reference used for calling the tool is different from the reference provided to VCF2Prot. These tests can be turned off by exporting the environment variable *NO_TEST* to minimize their run time cost especially for a repeated analysis of a well-studied dataset. Further, individual tests can be switched on by exporting the variable *RUN_SELECTED_TEST* followed by the test name, e.g. *INSPECT_INS_GEN*, which applies the above mentioned test to the generated instruction of each transcript.

### III Inspection and Validation of the Interpretation into Tasks

As discussed above, after translating the mutations observed in a transcript into *Instructions*, the sets of *Instructions* in a transcript are combined to generate a vector of *Tasks* donated as the IR of the sample-specific version of the transcript. Different reasons can cause incorrect generation of IR, for example, incorrect translation of mutations into *Instructions*, logical error in casting *Instructions*, or logical errors in interpreting instructions. These errors could be observed in edge-cases which are extremely rare and may have not been covered thoroughly during unit-testing and development. Hence, we developed two tests to validate the correctness of interpretation, first, expected length and second gapped sequences. In the first case, the expected length of the protein is computed given its set of *Instructions* and is compared to the length described by summing the length of each *Task*. In case the expected and computed length disagree, the interpretation is considered incorrect, and the transcript is skipped after printing a detailed error message to the user. The second test makes sure that the *Tasks* are resulting in an instruction without interruptions, here the vector of *Tasks*, i.e. the IR of the transcript is inspected for gaps according to equation 2,

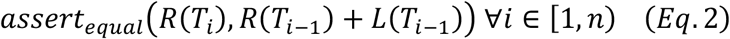

where *R* gives the position for a *Task (T*_*i*_) in the resulting protein, *L* gives the length of the *Task (T*_*i*_), finally, *n* is the number of *Tasks* (*T*s) in the IR, the *Tasks (T*_*i*_) are zero-indexed. Hence, in case the assertion of Eq.2 failed, the interpretation has failed as gaps are present in the resulting protein. Also, it is worth mentioning that this test is also used after concatenating the IR of each transcript to generate a haplotype-wide IR, to capture concatenation errors. By default, these tests are turned on, however, they can be turned off to increase the execution’s speed using the environmental flag NO_TEST. Also, specific test can be run by exporting the flag *RUN_SELECTED_TEST* as described above followed by the test name, e.g. *DEBUG_CPU_EXEC* to inspect the IR before executing it on the CPU.

## Results and Benchmarking

As stated above, available tools are written in different languages and are utilizing different input formats. Given that VCF2Prot specializes in analyzing genomic variants stored in a VCF file format, the most commonly used format to store genomic sequence information, we focused our benchmarking study on PrecisionProDB [6] as it supports the same input type. We used a python script to benchmark both tools against a VCF file containing an increasing number of samples ranging from 1 to 128 samples. As PrecisionProDB can only handle one sample at a time, a for-loop wrapper around the tool was used to measure and store the run time across all samples. As shown in **Fig. 2A**, VCF2Prot shows a massive improvement in the runtime and execution speed.

**Figure 2:**
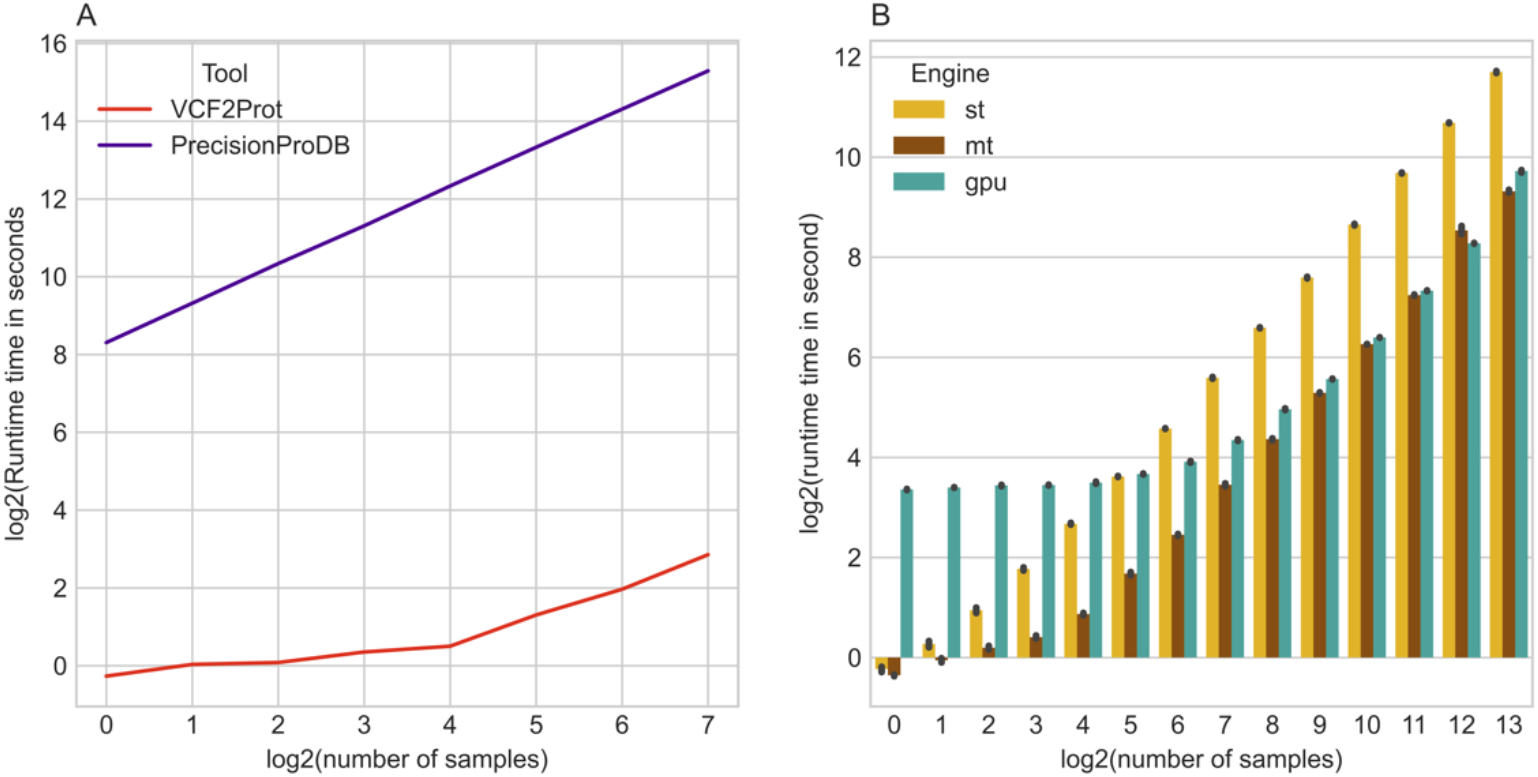
VCF2Prot performance benchmarking. A) The performance of VCF2Prot against PrecisionProDB using an increasing number of samples (logarithmic scale). Execution time is recorded in seconds and is shown on a logarithmic scale of base 2 (y-axis) where lower values indicate faster execution. For both VCF2prot and PrcisionProDB the multithreaded execution was enabled. B) A log-log plot for the execution time of VCF2Prot with an increasing number of samples in the input VCF file using different execution engines, namely, single-threaded (st), multi-threaded (mt) using a pool of 16 CPU threads, and graphical processing units (GPUs). Each dot represents the mean execution time over 12 different repetitions. For both A and B, the benchmarking was conducted on a GPU-node with 512GB of RAM, Twin Intel® Xeon® Gold 6134 CPU at 3.20GHz and using one Nvidia® Tesla V100-SXM2 32GB GPU.

Next, we were interested in benchmarking the scalability of VCF2Prot against an increasing number of samples. To this end we utilized a VCF file containing 99,254 variants observed across 17,138 individuals (***Suppl. Materials***). We benchmarked the three execution engines currently available in VCF2Prot, namely, single-threaded execution (st), multithreaded execution (mt), and GPU-execution (gpu) against an increasing number of samples in the VCF file (**Fig. 2B**). As seen in the figure, with a smaller sample size (n<64 =2^6^) using GPU increase the runtime. Meanwhile with growing sample sizes, this overhead of using GPUs becomes non-significant and we achieve a comparable performance between GPU-execution and multithreaded CPU-based execution. The multithreaded version is on average 4 folds faster that the single threaded execution (**Fig. 2B**).

## Discussion

The ability to generate sample-specific protein sequences is a cornerstone for a plethora of applications ranging from proteogenomics to neo-epitope discovery and peptidome-wide association studies. Different tools have been developed to address this task, nevertheless, most, if not all, of these tools have not been designed to handle large scale datasets, i.e. datasets with thousands of samples included. One of the challenges in benchmarking the output of different tools is the algorithm used for filtering and then calling the consequences of the genomic mutation. VCF2prot decouples the process of calling and identifying the alteration from the process of applying the alteration to generate the protein sequences. It relies on the state-of-the-art tool BCFTools/csq [9] for performing the consequence calling in a haplotype-aware manner and restricts itself to the subsequent processes of applying the genomic alterations to generate the sample-specific protein sequence. Thus, enabling the user to select and finetune the upstream processing pipeline, e.g., variant caller and/or consequence calling parameters.

Benchmarking VCF2Prot against the recently published state-of-the-art tool, PrecisionProDB shows a clear improvement in terms of execution speed and scalability where thousands of samples can be executed concurrently. Despite these advantages, the current version of VCF2Prot has still some limitations. First, it requires a phased and consequence called VCF file as an input which requires some preprocessing on the user-side and requires a large disk space to store the VCF file in comparison to the binary compressed BCF format. Second, VCF2Prot reads the whole file into memory and parses it. Although this improves the performance it might be challenging for users with large number of samples and a limited hardware infrastructure.

The simplicity of the internal representation (IR) generated by the SIR algorithm enables the generated vector of tasks to be executed on CPUs and GPUs. Nevertheless, benchmarking different execution engines showed a non-significant difference in the run-time of GPUs and multi-threaded CPU executions. Although it is possible to execute the vector of tasks on the GPU, it is not the rate-limiting step and hence, a future direction to improve the execution speed further would be the re-implementation of the preprocessing on the GPU. A second future direction is to enable VCF2Prot to parse different input format, e.g binary compressed and Tabix [10] indexed files along with support for writing the output results to an SQL database to improve down-stream task handling.

## Conclusion

SIR is an inherently parallel algorithm for generating personalized proteome sequences, which is a cornerstone for neoantigen-discovery and proteogenomics. The current implementation in VCF2Prot can process thousands of samples efficiently using CPUs and/or GPUs. The documented modular design of the implementation along with the permissive license shall help in developing other bioinformatics tools using the Rust programming language.

## Availability of data and materials

The source code of the library along with the precompiled executable are available at https://github.com/ikmb/vcf2prot, while the exome data used for characterizing the performance is available at: https://litdb.ikmb.uni-kiel.de/media/Regeneron_csq_1_min-ac5.bcf.gz

## Authors’ contribution

HE, MW, AF, FD, TLL designed and conceived the study. HE and MW developed the algorithm. HE implemented the tool in Rust. HE and MW wrote the manuscript. All authors read and approved the final manuscript.

## Acknowledgements

We are truly thankful for the help of Petr Danecek (Wellcome Trust Sanger Institute) for his prompt and responsive reaction in debugging *BCFtools/csq.* We would also like to thank Tim Steiert (Institute of Clinical Molecular Biology) for testing and validaing the tool.

## Supplementary Materials

### I Data Preparation and Consequence Calling

The example dataset comprised of whole exome-sequencing data for chromosome 1 of 17,138 samples. This dataset was assembled from different in-house WES projects. Sample preparation and sequencing as well as the sequence alignment, variant identification and genotype assignment were performed according to the protocol described in [11]. Additional genotyping was performed using the Global Screening Array (GSA), version 1.0 (Illumina) on the same samples. Genotype calling, extracting GSA genotyped data from intensity data files was performed with Illumina GenomeStudio v. 2.0 software using Cluster File GSAMD-24v1-0_20011747_A1. Standard genotype quality control was performed using BIGwas [12] with default parameters.

To perform haplotype aware consequence calling with *BCFTools/csq* [9] the data needed to be phased. To enable phasing with the phasing tool Eagle [13] the WES dataset and the GSA data needed to be merged in order to increase SNP coverage and enable phasing in otherwise sparsely covered regions of the exome data. As the WES data was called on genomic build GRCh38 and the GSA data on genomic build GRCh37, the GSA data was first lifted using LiftOver from UCSC Genome Browser [14]. All variants that mapped to non-classical chromosomes were removed (“--chr 1-24”), as well as duplicates (“--list-duplicate-vars suppress-first”, followed by “--exclude”) using PLINK, version 1.9 [15]. The variants in the WES and GSA data were split into overlapping variants and exclusive variants (i.e. present only in the WES or GSA data). The overlapping variants were merged into a temporary VCF file using PLINK. The GSA exclusive variants were exported to VCF with PLINK (--extract <snps> --recode vcf-iid) [15]. Finally, the GSA exclusive variants, exome exclusive variants and the overlap variants were merged with “bcftools concat” and afterwards sorted with “bcftools sort” [9].

Since multiallelic variants needed to be split into single entries for processing in Eagle, we applied “bcftools norm” to the merged and sorted VCF file [9]. Phasing was then conducted using Eagle, version 2.4.1, not imputing missing variants [13]. In case of a heterozygous genotype for an originally multiallelic position, phasing, in some instances, resulted in mapping of both variations on the same haplotype. Those cases were solved manually by randomly changing one variant to a valid combination. The multiallelic variants were then merged together using” bcftools norm −m +any” [9]. The variants were finally called using “bcftools csq -p a” (bcftools version: git_1f1e766)[9]) using the GFF flatfile from the ensemble reference for build GRCh38, release 100 [16].

## Supplementary Tables

**Table S1:**
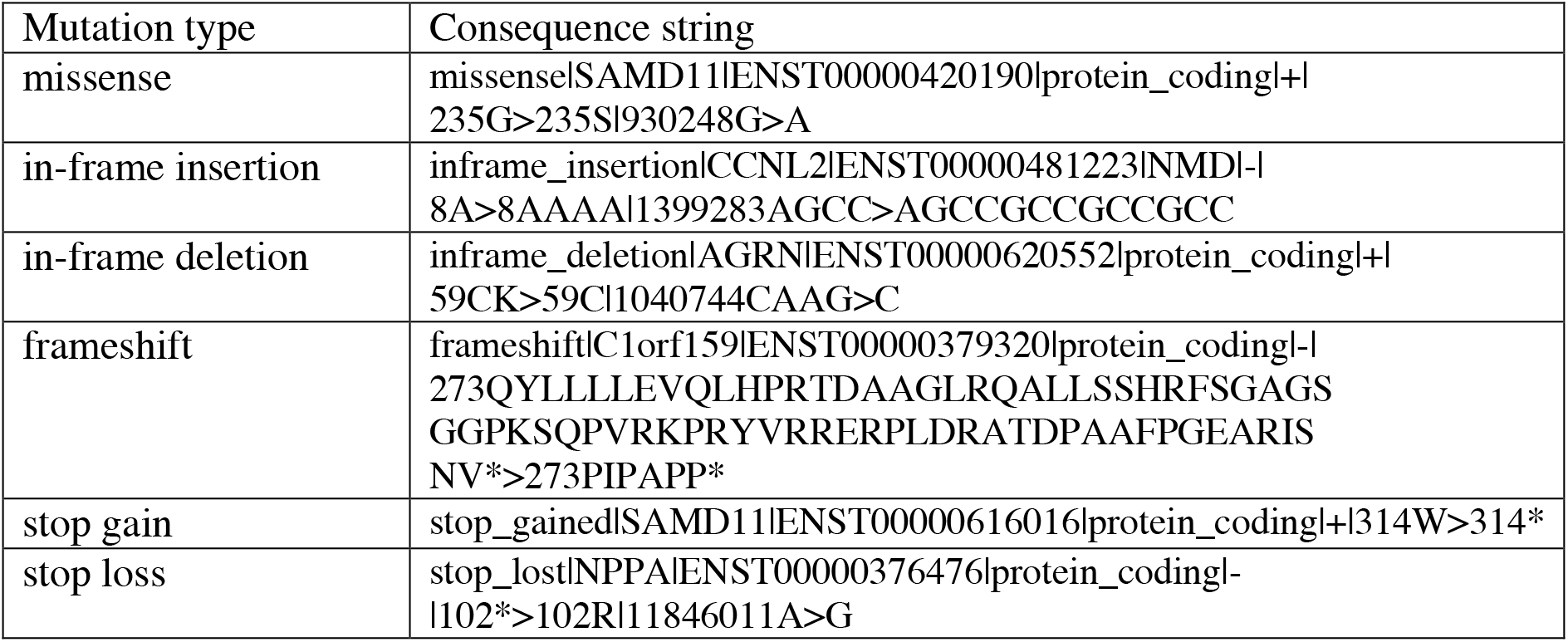
Examples of BCFtools/csq consequence string for most common mutational types. It is worth to mention that for stop-loss the follow up sequence remains unknown as it is not given by bcftools and hence only the sequence provided by the caller, i.e. BCFtools/csq is included into the resulting sequence.

**Table S2:** A summary of the supported genomic alteration types and the one letter code of each mutation. Phi is an introduced mutational type to represent a mutation or an alteration without an effect, e.g. a *missense upstream a frameshift or a stop-gain. Mutation names are given as described by BCFTools/csq [9].

| Mutation Name | Code |
| --- | --- |
| *frameshift | R |
| *frameshift&stop_retained | Q |
| *inframe_deletion | C |
| *inframe_insertion | J |
| *missense | N |
| *missense&inframe_altering | K |
| *stop_gained | X |
| *stop_gained&inframe_altering | A |
| frameshift | F |
| frameshift&start_lost | V |
| frameshift&stop_retained | B |
| inframe_deletion | D |
| inframe_deletion&stop_retained | P |
| inframe_insertion | I |
| inframe_insertion&stop_retained | Z |
| missense | M |
| missense&inframe_altering | Y |
| phi | E |
| start_lost | O |
| start_lost&splice_region | U |
| stop_gained | G |
| stop_gained&inframe_altering | T |
| stop_lost | L |
| stop_lost&frameshift | W |

## Supplementary Figures

**Fig. S1:**
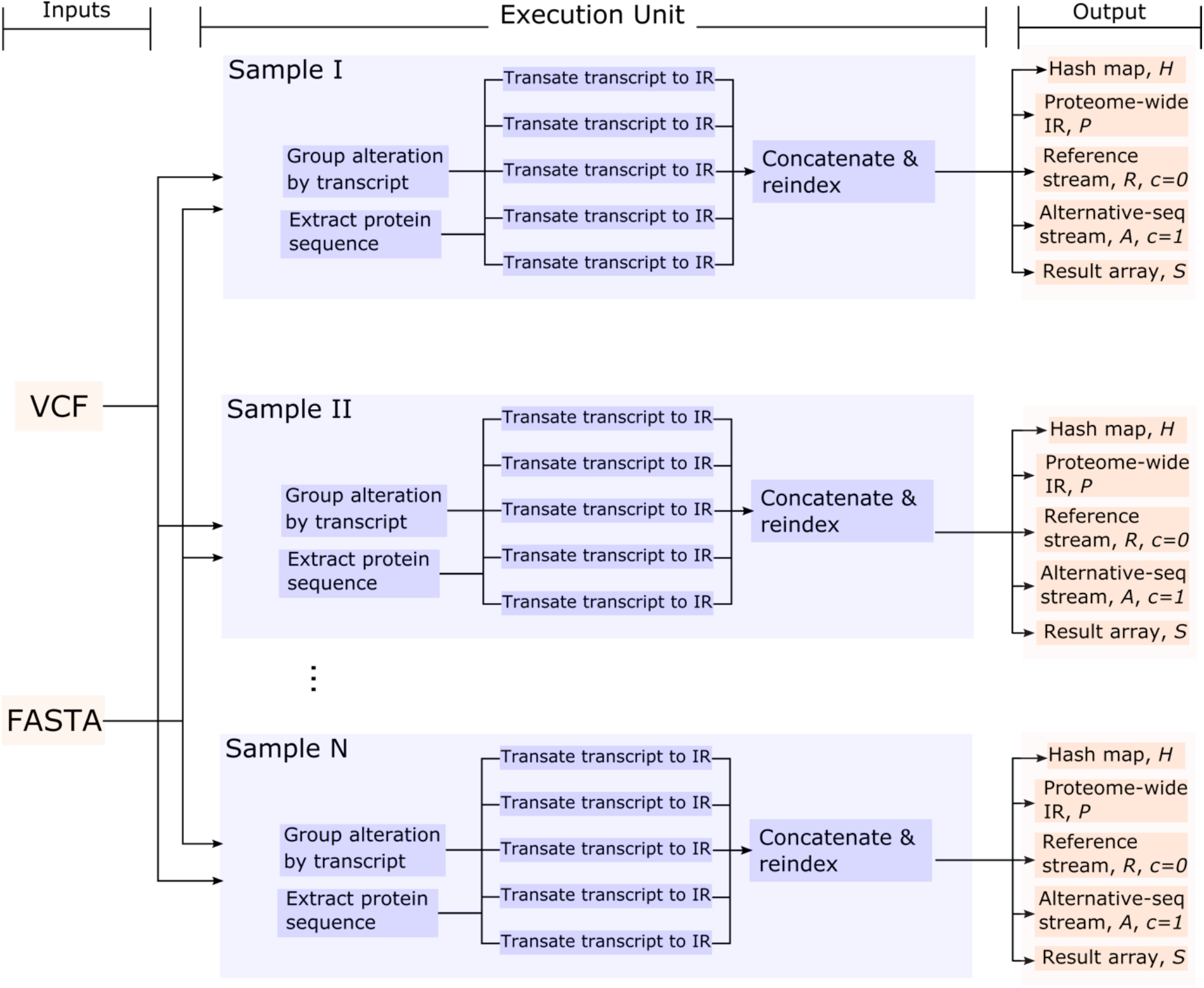
A graphical summary of sequence intermediate representation (SIR). The execution starts by reading an input VCF file and a FASTA file. Next samples are processed in parallel where for each sample the alterations are grouped by transcript-id and using a pool of threads each altered transcript is translated into an IR in parallel. Finally, the IR of all transcripts are concatenated and re-indexed for proteome-wide coordinates. It is also worth mentioning that the result array S, is an empty array that has been only allocated but will be filled when the IR is executed using any of the supported engines.

